# Disentangling principled and opportunistic motives for reacting to injustice: A genetically-informed exploration of justice sensitivity

**DOI:** 10.1101/2020.06.10.143925

**Authors:** Nikolai Haahjem Eftedal, Thomas Haarklau Kleppestø, Nikolai Olavi Czajkowski, Jennifer Sheehy-Skeffington, Espen Røysamb, Olav Vassend, Eivind Ystrom, Lotte Thomsen

## Abstract

Moral judgments may be driven by both principled and opportunistic motivations. Being morally principled is to consistently adhere to a single set of rules about morality and justice. Opportunistic morality rather involves selectively enforcing rules when they are beneficial to one’s interests. These two kinds of motivations sometimes pull in the same direction, other times not. Prior studies on moral motivations have mostly focused on principled morality. Opportunistic morality, along with its phenotypic and genetic correlates, remains largely unexamined. Here, utilizing a sample from the Norwegian Twin Registry, consisting of 312 monozygotic-and 298 dizygotic twin pairs (N = 1220), we measure people’s propensity to react to injustice as victims, observers, beneficiaries, and perpetrators of injustice, using the Justice Sensitivity scale. Our genetically informative sample allows a biometric modeling approach that provides increased stringency in inferring latent psychological traits. We find evidence for two substantially heritable traits explaining correlations between Justice Sensitivity facets, which we interpret as a *principled justice sensitivity* (h^2^ = .45) leading to increased sensitivity to injustices of all categories, and an *opportunistic justice sensitivity* (h^2^ = .69) associated with increased victim sensitivity and a decreased propensity to feel guilt from being a perpetrator. These heritable justice traits share a genetic substrate with broad strategies for cooperation (as measured by altruism and trust) and for selectively benefitting oneself over the adaptive interests of others (as measured by social dominance orientation and support for monopolizing territory and resources), and differ genetically and phenotypically from Big Five personality traits.

## Introduction

> *Many of our most serious conflicts are conflicts within ourselves. Those who suppose their judgements are always consistent are unreflective or dogmatic*
>
> — - John Rawls, Justice as Fairness: A Restatement, 2001

The question of how to ensure a truly fair system of justice may be as old as humanity itself. Having a shared standard of justice in a society can substantially improve the quality of life. But notions of justice can also be used as a tool by those in power to gain a disproportionate share of resources and privileges. On the interpersonal level as well, justice can be enforced strategically, so that a person gains more of the benefits of justice and incurs fewer of the costs. John Rawls (1971/2009) argued that, ideally, systems of justice should be shaped from behind a *veil of ignorance*, where decision makers do not know their own standing in society, and are thus unable to make rules that unfairly benefit people like them. Similar sentiments have also been put forward by others: Adam Smith claimed justice should be decided by *impartial spectators* (1759/2010); David Hume valued the *common point of view* (1739/2012); and Roderick Firth’s *ideal observer* is completely neutral (1952).

In Rawls’ scenario, the corrupting influence of self-interest on justice is mitigated through fully aligning self-interest with the shared interests of everyone. A person behind a veil of ignorance is made to weigh equally the outcomes for all members of society, because they do not know where they will end up. This then leaves open the question of whether people can ever have truly impartial and principled motivations towards justice, rather than merely selfish motivations that happen to align with the common good. Do people value justice for its own sake, beyond just pursuing a good deal for themselves and those they care about when negotiating a social contract?

Classical theories on the evolution of morality suggest that people indeed have such conflicting moral motivations. Moral sentiments are here thought to originate from the dynamic relation between cooperation and defection (cf. Wilson, 1998; Trivers, 1971). The principled employment of common moral rules for mutually beneficial cooperation (e.g., for reciprocity) might yield adaptive advantage in the context of a highly social species (cf. Boyd & Richerson, 2005; Henrich, 2017) but is only possible to sustain insofar as cooperators are able to discriminate and deter defectors from reaping the benefits of cooperation without contributing to its costs (cf. Trivers, 1971). Conversely, defectors will benefit from not being found out and punished. Evolutionary theory suggests that this will create an evolutionary arms race of adaptations, counter-adaptations, and counter-counter-adaptations for cooperative and defecting strategies regarding moral behavior—and their genetic substrates—within the species (Dawkins & Krebs, 1979; Wilson, 1998).

An emerging literature in moral psychology provides some empirical support for these notions from evolutionary theory (e.g., Simler & Hanson, 2017; Mercier & Sperber, 2011). On the one hand, people can feel genuine commitments to moral standards, such that they are motivated to enforce these standards consistently, regardless of whether they stand to gain or lose in the current context (e.g., Sinnott-Armstrong & Miller, 2017). On the other hand, people sometimes employ moral rules opportunistically – but in a manner allowing them to plausibly deny that they do so. For instance, the appropriate level of blame for a given moral violation can be adjusted up or down depending on whether you side with the violated or the violator (e.g., Eftedal & Thomsen, 2020), and the criteria for what is a moral violation in the first place could also be shifted, from one situation to the next (Clark, Chen, & Ditto, 2015). In other words, people generally endorse universal moral standards, but they systematically shift these standards to better serve their interests.

In his work on self-deception, Trivers argues that self-serving calculations likely happen mostly sub-consciously because it is easier to convince others of your value as a trustworthy, cooperative social partner if you have already convinced yourself (Trivers, 2011). Applying this logic to the domain of justice, the conscious mind would need to keep seeing itself as impartial and just, so as to better be able to convince others. If opportunistic morality is indeed mostly sub-conscious, this will make it harder to tell from the outside the extents to which moral motivations are principled or opportunistic. So too when using scale measures from moral psychology. As we will argue, however, these two kinds of motivations can be teased apart by looking at patterns of judgments across different situations. Specifically, it is informative to look at how judgments change when one’s perspective to injustice changes.

In the present study, we investigate this perspective on moral judgment using the Justice Sensitivity (JS) scale (Schmitt, Gollwitzer, Maes, & Arbach, 2005; Baumert & Schmitt, 2016). This scale measures how readily someone perceives- and reacts to injustice in the world around them.

Importantly, the scale has four facets, reflecting how the propensity to perceive and react negatively to injustice can vary depending on the perspective from which the injustice is experienced. The four perspectives are those of a *Victim*, an *Observer*, a *Beneficiary*, and a *Perpetrator*. The *Victim* facet represents sensitivity to injustice of which you are yourself a victim. *Observer* is sensitivity to injustice you observe as a bystander, without being directly involved. *Beneficiary* represents a tendency to feel guilty about passively benefiting from injustice. *Perpetrator* is the propensity to see one’s own behavior as constituting injustice towards others, and to then feel shame and guilt.

A principled motivation to uphold justice for its own sake would influence sensitivity to injustice equally across these four perspectives. A violation of justice is then equally bad regardless of whether you are a victim, an observer, a beneficiary, or a perpetrator. Thus, to the extent that people vary in having this kind of principled motivation for preserving justice, we predict a *principled justice sensitivity* factor to exist; a tendency to object to injustice regardless of one’s perspective to it.

Conversely, with an opportunistic motivation to use justice norms in service of other goals, the perspective to injustice can indeed impact one’s sensitivity. We thus predict an *opportunistic justice sensitivity* factor, reflecting a tendency to object more strongly to injustice that harms you or those you care about, and less strongly to injustice you perpetrate and/or benefit from.

Our large genetically-informative sample of mono- and dizygotic twin pairs allows a more comprehensive approach to identifying these kinds of latent traits (Franic et al., 2013). And it enables us to explore the extents to which principled and opportunistic morality are genetically or environmentally formed.

Insofar as principled and opportunistic justice sensitivity reflect general cooperative versus defecting social strategies, they may also share genetic substrates with other general cooperative strategies, such as those related to altruism and trust, as well as with broad strategies to maximize personal/in-group gain at the expense of others, such as dominance motives and support for policies to monopolize territory and resources. Indeed, recent evidence suggests that such motives for dominance versus equality form are grounded in a common, partly heritable genetic substrate (Kleppestø et al, 2019). Here, we test if this common genetic substrate also extends to the opportunistic employment of the moral rules of justice.

### Principled Justice Sensitivity

To have a principled moral motivation for justice is to see justice as valuable in itself, regardless of who stands to lose or benefit from it in a given situation. David Hume thought such motivations exist, proposing that we have a moral sense which produces intuitions from a purely impartial perspective; “the common point of view”. These intuitions then compete with all our other concerns before they influence behavior. Recent work in moral psychology supports Hume’s view that we can indeed have genuine commitments to moral standards (see e.g. Sinnott-Armstrong & Miller, 2017). Being seen by one’s social group as impartial and committed to moral rules is valuable from a game theoretic standpoint (Schelling, 1960), as this increases others’ expected returns from mutually beneficial cooperation. Consistent with this, the preservation of a narrative of moral consistency to self and others could be a basic psychological need (Prentice et al., 2019; Vonasch et al., 2017). If people indeed experience principled moral motivations, and if they do so to varying degrees, then this should contribute to positive correlations between the four facets of the Justice Sensitivity scale. An increased concern with justice for its own sake would lead to higher sensitivity to all injustice, regardless of whether one is a victim, observer, beneficiary, or perpetrator. We label this hypothesized common factor *principled justice sensitivity* (Principled-JS).

While each perspective on justice sensitivity might be influenced in the same way by such a morally-principled trait, they also differ, for example, in the emotional responses they tend to elicit. Being a victim or observer of injustice typically elicits anger and indignation, while being a beneficiary or perpetrator rather leads to shame and guilt (Mikula, 1994; Mikula, Petri, & Tanzer, 1990). Furthermore, being a victim or perpetrator can be distinguished from being an observer or beneficiary in that the former involve a higher degree of personal involvement, leading to more intense reactions. The four-facet structure of the JS scale reflects these differences and has been supported by several studies, both for a long version of the scale with ten items per facet (Schmitt et al., 2005; Schmitt, Baumert, Gollwitzer, & Maes, 2010), and for a short version with two items per facet (Baumert et al., 2014). Nevertheless, the empirical record suggests that the four JS facets do tend to correlate substantially, mostly at around r = .50 (Baumert & Schmitt, 2016), consistent with the existence of a general trait of principled justice sensitivity. While models combining pairs of JS facets have been shown to produce worse fit (Schmitt, Baumert, Gollwitzer, & Maes, 2010), it might still be the case that a principled justice sensitivity exists *in addition to* specific sensitivity to being the victim, perpetrator, observer, or beneficiary of injustice. This would be represented by a single factor model (Figure 1.a.) which also allows for influences that are specific to each facet of the JS scale, as represented by the correlated errors on items belonging to the same facet.

**Figure 1.**
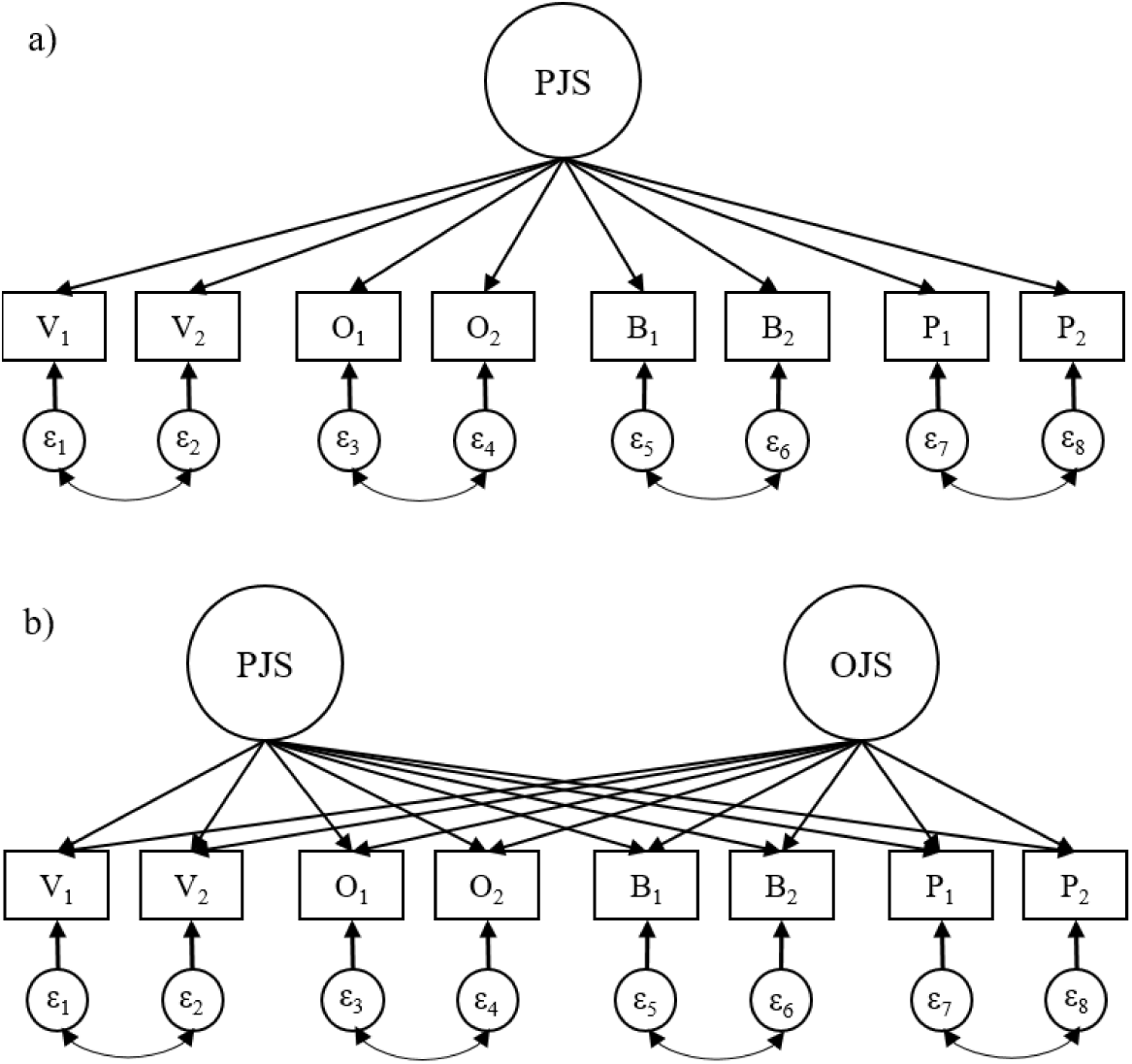
Illustrations of factor models to explain correlations between JS-facets. **Note.** V, O, B, and P represent the JS facets of Victim, Observer, Beneficiary, and Perpetrator sensitivity, respectively, with numbers distinguishing between different items in the scale from the same facet; PJS is Principled Justice Sensitivity; OJS is Opportunistic Justice Sensitivity. a) Model with a Principled-JS trait influencing all facets; c) Model with separate Principled-JSand Opportunistic-JS traits influencing the JS-facets.

However, this principled justice sensitivity trait fails to explain an interesting pattern in how justice sensitivities correlate: victim and perpetrator sensitivity tend to correlate only quite modestly with each other, even as they both correlate substantially with both observer and beneficiary sensitivity (see e.g. Baumert et al., 2014; Schmitt et al., 2010). This pattern suggests an additional factor could be at play, which pulls victim and perpetrator sensitivity in opposite directions (Figure 1.b.).

### Morally Opportunistic Justice Sensitivity

The idea that people may use moral norms about justice in instrumental, opportunistic ways is supported by an emerging literature in moral psychology and motivated reasoning (e.g., Simler & Hanson, 2017; Mercier & Sperber, 2011). For instance, in judging the severity of violations of norms related to freedom of speech, people assessed restrictions of speech they supported as more blameworthy than the same restrictions imposed on speech they opposed – even while explicitly denying that speech content should play any part in such judgments (Eftedal & Thomsen, 2020). Such moral selectivity, too, appears to be a trait which people possess to varying degrees. In the context of Justice Sensitivity, we will label this trait of being self-servingly selective about morality as *Opportunistic Justice Sensitivity* (Opportunistic-JS).

Evidence that opportunistic moral selectivity (or at least its inverse) may be an enduring personality trait is found in studies of the Honesty/Humility-dimension of the HEXACO personality inventory, tapping trustworthiness, modesty, and a lack of greed (Ashton et al, 2004; Ashton & Lee, 2009), and of the Morality facet of Agreeableness in the IPIP-NEO, which reflects an aversion to using cheating and deception in order to get ahead (Goldberg et al., 2006). Indeed, such measures have been found to be associated with responses to the JS scale: Baumert, Schlösser, and Schmitt (2014) found that the honesty/humility factor correlated negatively with victim sensitivity but positively with the sensitivity to being the beneficiary, observer, and, most substantially so, the perpetrator of injustice.

Presumably, an opportunistic selectivity in using moral rules would increase victim sensitivity and decrease beneficiary and perpetrator sensitivity. High victim sensitivity can lead to increased compensation in terms of resources and privileges, while decreased beneficiary and perpetrator sensitivity implies a decreased willingness to compensate others. Thus, opportunistic justice sensitivity maximizes material gains and minimizes losses from being sensitive to injustice, at least in the short term (until others catch on to one’s true motivations). Consistent with this proposal, victim sensitivity has been found to predict selfish behavior across several kinds of economic games (Fetchenhauer & Huang, 2004; Gollwitzer et al., 2009), and to correlate positively with Machiavellianism, neuroticism, and jealousy (Schmitt et al., 2005). Furthermore, the people who are most opportunistic or Machiavellian in their use of moral rules have been found to be the least prone to feelings of shame and guilt, which are in turn the main emotions associated with sensitivity to being a perpetrator of injustice (Montebarocci et al., 2004). Indeed, opportunistic justice sensitivity may especially reduce sensitivity to being the perpetrator of injustice, because expectations for you to personally compensate others are stronger if you have perpetrated an injustice yourself, as opposed to simply observing or passively benefitting from injustice (Alicke, 2000). The possibility that opportunistic justice sensitivity might be another latent trait, in addition to principled justice sensitivity, which underlies the relationships between sensitivity to being the victim, perpetrator, observer, and beneficiary of injustice, is represented by a two-factor model (Figure 1.b). Here, all items are specified to load on both a principled and an opportunistic latent factor (in addition to correlated errors for items belonging to the same JS-facet).

In summary, there is good reason to expect that in addition to Principled Justice Sensitivity, Opportunistic Justice Sensitivity will be a second latent trait underlying the pattern of correlations between sensitivities to being the victim, perpetrator, observer, or beneficiary of injustice. In contrast to a principled motivation, which is predicted to have a uniform effect across these four JS facets, we predict that an instrumental, opportunistic motivation for justice will increase sensitivity to being a victim, but decrease sensitivity to being a beneficiary or a perpetrator. The oft-reported finding of a comparatively low correlation between victim and perpetrator sensitivity might then reflect a complex relationship where they are both pulled in the same direction by principled motivations for justice but also (less strongly) in opposite directions by opportunistic motivations.

### Exploring justice concerns with gentically informative data

Traditional approaches to identifying latent psychological traits, such as confirmatory factor analyses, are solely reliant on phenotypic correlations. Such approaches risk underestimating the complexity of the explanation for why the observed variables correlate (e.g., Franić, et al., 2013). A pattern of correlations that supports inferring a small number of latent traits in a standard factor analytic (phenotypic) approach could arise from a situation involving a much larger number of separate influences (see e.g., Thomson, 1916). This shortcoming has led to the development of biometric approaches to identifying latent traits (e.g., Franić, et al., 2013), which involve using data from twins reared together to compare the patterns of genetic and environmental influences on phenotypic scores from across a set of items. The logic is that in a situation where latent psychological traits cause individual items to correlate, then genetic and environmental patterns of correlations between items should be highly similar. All genetic and environmental influences on phenotypic correlations should then work via the same latent trait.

We use this biometric approach to investigate latent traits underlying sensitivity to injustice across the perspectives of the victim, the observer, the beneficiary, and the perpetrator. In addition to allowing increased confidence in inferring the presence of latent traits, our approach also enables exploring the heritability of traits we identify.

Our prediction of separate principled and opportunistic justice sensitivities corresponds a classical theoretical proposal that the biology of morality is undergirded by an intra-species evolutionary arms-race of adaptations and counter-adaptations for social manipulation between cooperative and defecting strategies for social manipulation (cf. Dawkins, 1979; Wilson, 1998; Trivers, 1971, 2011). Principles of balancing selection lead to the expectation of heritable variation in those behavioural orientations whose adaptive value depends on the nature of the ecological context, where the latter includes the distribution of other traits in a population (cf. Arslan & Penke, 2015). Whether or not to endorse moral principles, and whether one applies them impartially, arguably fits this description of a behavioural orientation with systematically variable adaptive value.

Our analyses will also bear on debates in the fields of personality and moral psychology regarding the nature and origins of moral character. There is research suggesting that situational and structural factors impact attitudes and beliefs about justice (e.g. Wijn & van den Bos, 2010). On one hand, this might suggest that shared environmental factors among twins, such as the socioeconomic status and parenting, will play a more prominent role in explaining variability in such moral justice concerns than is typically found for other psychological traits. Shared environmental factors like these typically account for less than 15% of variance in psychological traits, while genes typically account for between 30% and 60% of variability (Vukasović & Bratko, 2015). On the other hand, several other pro- and anti-social traits that are relevant for cooperative versus opportunistic morality have been found to be heritable, including dark triad characteristics of Machiavellianism (Vernon et al., 2008), psychopathy (Tuvblad et al., 2014), dispositional empathy (Davis, Luce, Kraus, 1994), altruism (Rushton et al, 1986) and trust (Cesarini et al, 2008). This suggests that moral sensitivity to injustice may also be substantially heritable. It is thus valuable to use a genetically-informative sample to estimate the relative contributions of genes, the shared environment, and other influences to explain variability in justice sensitivity.

Finally, biometric modeling can shed light on justice sensitivity’s relationship to, and distinctness from, related psychological traits. One pressing question concerns whether any such heritable genetic substrate for justice sensitivity is distinct from, or simply reduces to, Big Five personality traits (e.g., agreeableness) (John & Srivastava, 1999) that are phenotypically correlated with justice sensitivity (Schmitt et al., 2010), and are themselves highly heritable (Vukasović & Bratko, 2015). This addresses the general theoretical question of whether moral character and values are distinct from the temperament and action tendencies which personality inventories typically tap (Cawley III, Martin, & Johnson, 2000). Also important to analyse is whether any such distinct, heritable strategies for principled versus opportunistic justice sensitivity share common genetic substrates with other psychological traits that serve to sustain cooperation for the greater common good—such as altruism (Rushton et al, 1986; Wang & Lu, 2018) and trust (e.g., Cesarini et al, 2008)—versus serve to maximize adaptive fitness gains over others, such as social dominance and the monopolization of territory and resources (Kleppestø et al, 2019). This possibility can be tested with our genetically informative sample.

### The Present Research

We run and compare a series of biometric factor models to test if morally principled and opportunistic strategies with distinct, heritable substrates undergird sensitivity to being the victim, perpetrator, observer, and beneficiary of injustice. We compare genetically-informative Common or Independent Pathway (CP and IP) models, and we also run a standard phenotypic confirmatory factor analysis for reference. We also use factor scores to partition the multivariate variance into genetic and environmental correlations, enabling us to investigate how principled versus opportunistic justice sensitivity relate to Big Five personality traits, altruism, trust, social dominance orientation, and support for policies that monopolize territory and resources.

## Materials and Methods

### Sample

The twins in our sample were recruited from the Norwegian Twin Registry (NTR), which consists of several cohorts of twins. Our cohort consisted only of same-sex twins, born between 1945 and 1960. The questionnaire was filled out in 2016. The mean age was 65.16 (SD=4.49). For complete pairs with valid scores on our scales of interest, our sample consisted of 121 male monozygotic twin pairs, 124 male dizygotic pairs, 191 female monozygotic pairs, and 174 female dizygotic pairs. This yielded a total of 610 twin pairs, or 1220 subjects. Zygosity was determined by a questionnaire shown to correctly classify >97% of twins (Magnus, Berg, & Nance, 1983).

### Measures

Among the measures in the questionnaire, we include the following measures in our analyses:

#### Justice Sensitivity (JS)

We used the short version of the Justice Sensitivity scale, which has a total of eight items; two items each for *Victim-, Observer-, Beneficiary-*, and *Perpetrator-*sensitivity. For example, the item “It makes me angry when others are undeservingly better off than me” is part of the Victim-facet. Agreement to each item was indicated using a 7-point Likert scale (1 = strongly disagree, to 7 = strongly agree). Cronbach’s α for the scale as a whole was .77.

#### Social Dominance Orientation (SDO)

Social dominance orientation was measured with the SDO_6_ (Pratto et al., 1994), a 16-item scale that reflects one’s preferences for the unequal distribution of power and resources between groups in society (α = .85).

#### Interpersonal Trust

Our measure of interpersonal trust is from the European Social Survey (www.europeansocialsurvey.org). It consists of three items, and reflects whether respondents tend to see other people as generally trustworthy (α = .82).

#### Altruism

The Self-Report Altruism scale measures the self-reported frequency of a set of five selfless behaviors, such as donating blood or money (Rushton, Chrisjohn, & Fekken, 1981) (α = .48).

#### The Big Five Inventory (BFI)

The BFI is a well-established taxonomy of personality, with the five dimensions of *Openness to experience* (α = .78), *Conscientiousness* (α = .73), *Extraversion* (α = .80), *Agreeableness* (α = .72), and *Neuroticism* (α = .83). Our survey included the BFI-44 (John & Srivastava, 1999), which is a shortened 44-item version of the full BFI scale.

#### Political attitudes about immigration and foreign aid

Our survey included four questions about attitudes towards immigration policy and foreign aid. Responses to these questions correlated highly, and so we combined them into a single scale (α = .77) where high scores correspond to less support for immigration and foreign aid.

### Analyses

#### Twin models

The classical twin design allows partitioning the variation of a trait into three components: *A*, C, and *E*. A is additive genetic influences, C is shared-environmental influences, and E is unique-environmental influences. Genetic effects cause monozygotic (MZ) twins to be more similar to each other than dizygotic (DZ) twins are; shared-environmental effects are everything that makes twins similar to each other except genetics; and unique-environmental effects are everything that makes twins different from each other except genetics. Thus, in addition to individual experiences, the unique environment also contains measurement error as well as the intrinsic molecular stochasticity in the process of building a phenotype from a genotype (Tikhodeyev & Shcherbakova, 2019).

We used structural equation modeling to model the covariances of the responses of twins in terms of additive genetic effects, shared-environment effects, and unique-environment effects (see e.g. Neale, 2005). These models were fitted by full-information maximum likelihood analysis using OpenMx (Boker et al., 2011). As we focus here on general aspects of Justice Sensitivity rather than sex differences, sex was regressed out of the justice sensitivity scores, and standardized residuals were used in subsequent twin analyses (McGue & Bouchard, 1984).

#### Common- and independent pathway biometric models

As an elaboration on standard factor analytic models with latent factors, twin data allow fitting models with separate A, C, and E components. These are called Independent Pathway models (“IP model”; see figure 2a), and a set of separate A, C, and E components constitute a single “IP-factor”. Such a model then partitions the sources of covariances into A, C and E.

**Figure 2.**
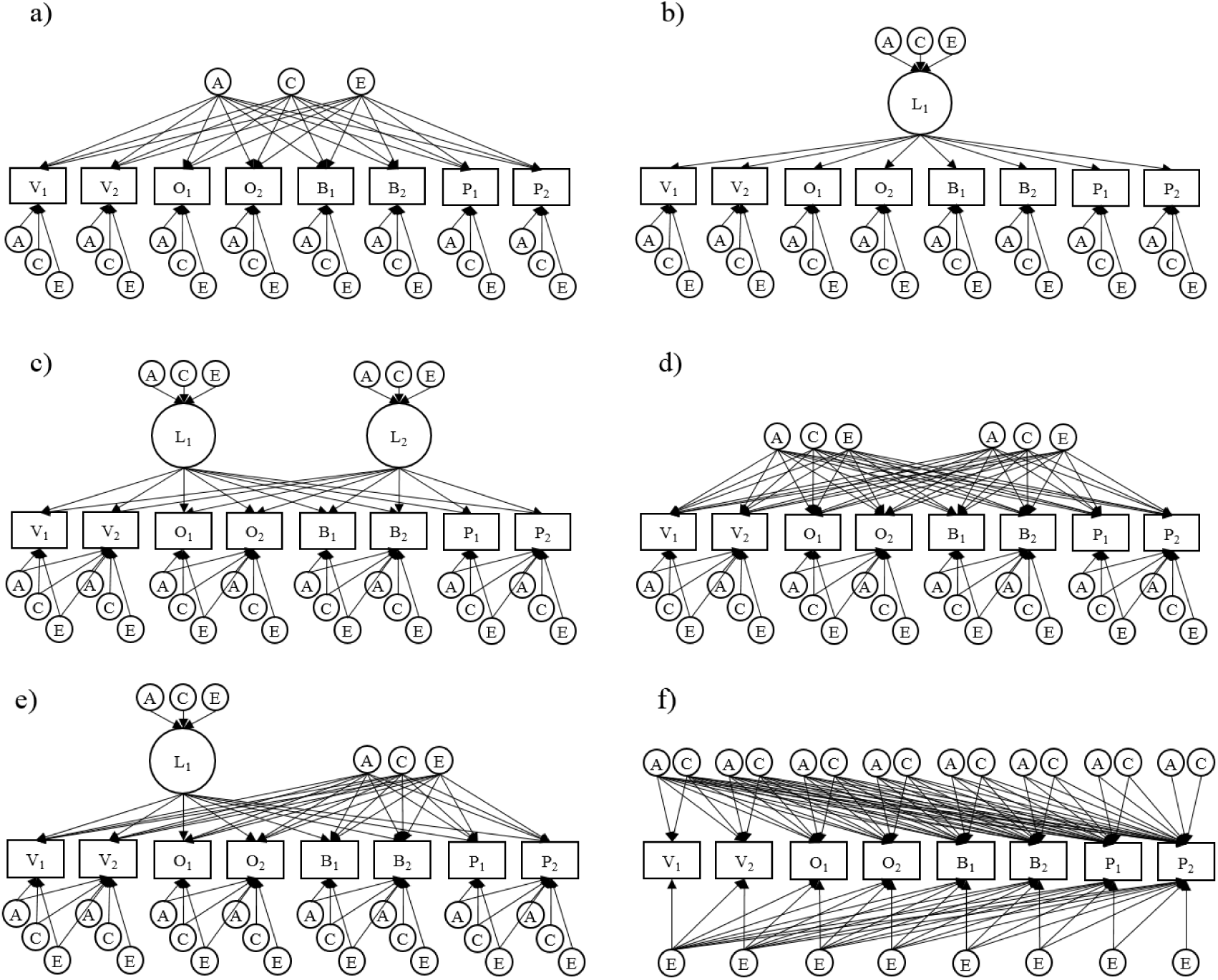
Illustrations of multivariate twin models. *Note*. a) 1-factor IP-model; b) 1-factor CP-model; c) 2-factor CP-model allowing items from the same JS-facet to share variance beyond what is accounted for by the common factor (the models in d and e also allow for this); d) 2-factor IP-model e) hybrid model with one CP-factor and one IP-factor; f) Cholesky-model, working as our baseline for comparisons.

If, in fact, the true reason for the covariances between observations is the existence of a single latent trait, then the patterns of loadings (i.e. the relative magnitudes of loadings) from the A, C, and E components should be equal. After all, in this situation, genetic and environmental inputs could only affect covariances through first affecting the latent trait, which has a specific pattern of loadings on the observed variables. When adding such a latent trait to the model, and specifying that all influences on covariances of items must occur through first affecting this trait, the model becomes a Common Pathway model (“CP model”; figure 2b). A CP model is then equivalent to an IP model where loading patterns from the A, C, and E components are constrained to be equal. Adding this constraint reduces the number of estimated parameters, and thus will lead to improvements in parsimony-corrected fit indices if covariances are indeed explained by a single latent variable. Conversely, the more divergence there is from this situation of a single latent trait, the more genetic- and environmental loading patterns will tend to differ (for more details, see e.g. Franic et al., 2013). IP models would then become superior to CP models. If an IP model is better than a CP model in a situation where phenotypic models suggest a one-factor solution, this suggests that the real reasons for the covariances are in fact more complex than just the single factor from the phenotypic model.

For cases with more than one latent factor, such as in our hypothesized scenario with both Principled-JS and Opportunistic-JS traits, it is possible to compare CP and IP models with more than one factor. A CP model would then correspond to there being several *sets* of genetic and environmental factors with the same patterns of loadings; one set for each latent trait (figure 2c). IP models are equal to these models, except that they again lack the constraint of equal loading patterns; for each latent trait, separate A, C, and E factors are estimated (figure 2d). In a situation with two factors, it is possible that one them is a genuine latent trait, while the other one is not. Such a situation can be represented by a CP-IP hybrid model, with one CP-factor constrained to have equal loading patterns, and one IP-factor without this constraint. (figure 2e). We compare models with either one, two, or three factors, in CP-, IP-, and CP-IP hybrid versions. Models with two or less factors represent the existence of at least one of the hypothesized Principled-JS- and Opportunistic-JS traits. Models with three factors allow for the existence of additional, unforeseen traits also contributing to correlations between JS-facets. IP-models with multiple factors can only be fitted with orthogonal factor rotations, so we only investigate orthogonal factor rotations for CP-models as well, for cleaner comparisons.

Variance for each item that is unexplained by the latent factors is divided into separate A, C, and E components. To account for how items belonging to the same JS-facet share variance beyond that explained by latent factors, our models allow the variance components for the first item in a pair to also load on the second item in the pair.

As a baseline for comparisons, we use a model with a standard Cholesky decomposition of the variance shared between items (figure 2f). This is a type of model that simply allows all the observed variables to have correlated genetic and/or environmental influences, without specifying any broader latent factors as the sources of these correlations. It is a highly flexible model that can accommodate most kinds of explanations for why the JS-facets correlate, including situations without a simple structure with a small number of factors.

In addition to the full ACE versions of the models, we estimate AE and CE versions where C or A components are removed. An IP-factor in an AE model then consists of just an A-factor and an E-factor, and likewise for CE models. Since our approach is still quite far from exploring the full space of possible models, we also investigate if our best-fitting model can be further improved by removing or adding single A, C, or E components.

Since the models we are comparing are not nested within the baseline Cholesky model, we use Akaike’s Information Criterion (AIC; Akaike, 1974) to adjudicate between them.

For completeness, we also conduct a traditional phenotypic Confirmatory Factor Analysis on our data, assessing for the same set of two latent traits.

## Results

### Descriptives

Descriptive statistics for all measures and twin correlations are given in Table 1.

**Table 1.** Descriptive statistics and twin correlations.

| JS-facet | r | Mean (SD) | Twin correlations (95% CI) |  |
| --- | --- | --- | --- | --- |
|  |  |  | MZ | DZ |
| Victim | .70 | 2.69 (1.34) | .35 (.28, .41) | .17 (.10, .24) |
| Observer | .71 | 4.32 (1.42) | .29 (.22, .36) | .12 (.04, .19) |
| Beneficiary | .80 | 3.00 (1.50) | .30 (.23, .37) | .10 (.02, .17) |
| Perpetrator | .81 | 5.25 (1.81) | .26 (.19, .33) | .16 (.08, .24) |

*Note.* r = correlation between the two items for each facet.

Table 2 shows the correlations between facets. Here, it is noteworthy that the correlation between Victim and Perpetrator is even lower than in other contexts, at .08.

**Table 2.** Phenotypic correlation matrix for the four JS-facets (with 95% CIs).

|  | JS-Victim | JS-Observer | JS-Beneficiary |
| --- | --- | --- | --- |
| JS-Observer | .34 (.30, .38) |  |  |
| JS-Beneficiary | .38 (.34, .42) | .47 (.44, .50) |  |
| JS-Perpetrator | .08 (.03, .12) | .32 (.27, .36) | .29 (.25, .33) |

*Note.* All correlations are significant at $p < .01$ .

### Confirmatory Factor Analysis

Exploring whether correlations between JS-facets are best described by one or two latent factors first in a standard CFA-framework, we fitted the two models shown in Figure 1. All models were fitted with standard errors that were correlated for each twin pair, to account for non-independencies of observations. We also fitted these models without the correlations between residuals from items belonging to the same facet. Model fit deteriorated substantially when these correlations were removed. See Table 3 for statistics for the different models.

**Table 3.** Comparisons for phenotypic CFA.

| Model | EP | AIC | RMSEA | CFI |
| --- | --- | --- | --- | --- |
| 1-factor no-cor | 16 | 40991.48 | .198 | .58 |
| 1-factor cor | 20 | 39020.09 | .062 | .97 |
| 2-factor no-cor | 24 | 39850.77 | .160 | .80 |
| **2-factor cor** | **28** | **38884.62** | **.033** | **.99** |

Note. "cor" and "no-cor" signify whether a model is fitted with- or without correlations between residuals for items belonging to the same facet, respectively; EP = number of estimated parameters; AIC = Akaike's Information Criterion; RMSEA = Root Mean Squared Error of Approximation; CFI = Confirmatory Fit Index. Best fitting model indicated in bold.

The model with two latent factors (figure 1b) provided better fit than the model with only one such factor (figure 1a). RMSEA for this model was 0.033 and CFI was 0.99, meaning that it fit the data adequately.

The full model with parameter values is shown in Figure 2. The first factor aligns with our proposed Principled-JS trait, with substantial positive loadings on all JS-items. The second factor does also align with Opportunistic-JS to some extent, in that it has positive loadings on JS-Victim and negative loadings on JS-Perpetrator. Less expectedly, loadings on JS-Beneficiary are positive, and loadings on the second item in each pair from the same JS-facets are consistently more positive than for the first item in the pair (particularly so for JS-Observer).

**Figure 2.**
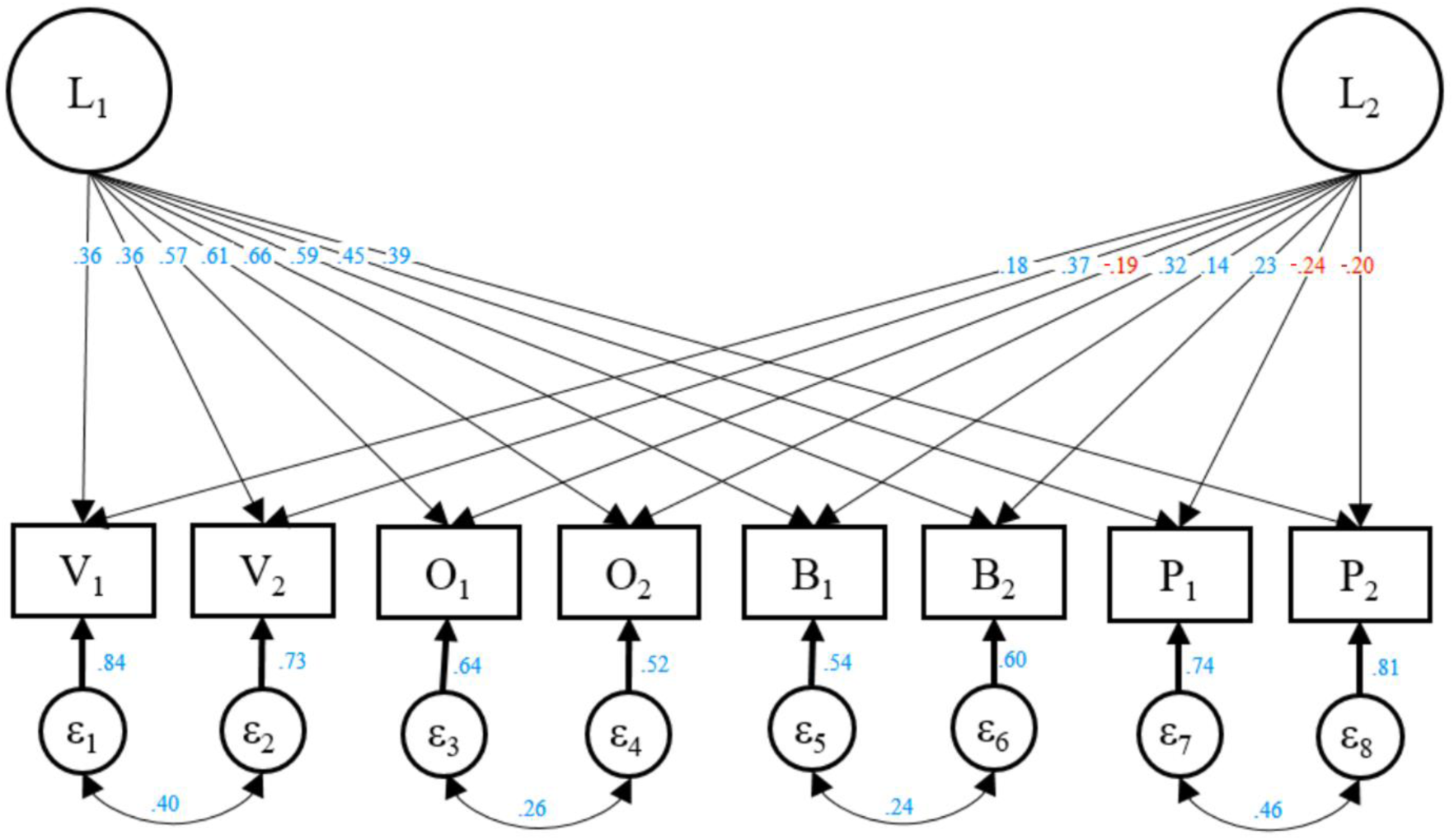
Best fitting model from phenotypic CFA. *Note.* Illustration of our best fitting model from phenotypic CFA. Blue numbers are positive and red numbers are negative.

### Biometric factor models

We compare CP, IP, and CP-IP hybrid models with one, two, or three factors (see Figure 2 for illustrations). As a baseline model for comparisons we use a full Cholesky decomposition of variance. The fit statistics for these models are detailed in Table 3. Pure AE models (with all C’s taken out) were consistently superior to models removing only a subset of the C’s. So, for clarity, we present statistics only for pure AE models here (except the baseline model, “Cholesky ACE”). Similarly, allowing items from the same JS-facet to share variance beyond that accounted for by common factors was also consistently preferable, so all models in the table have this property as well.

Our best-fitting model from this initial comparison was a 2-factor CP-IP hybrid model (CP1_IP1). As large parts of the space of possible models is still unexplored, we tested adjustments to this model involving the addition of a single variance component. This new component either comes in the form of an additonal pure A or E factor, or it can be added to the existing A or E factors making up the IP-factor in the model so that they become CP-factors. (Models involving the subtraction of variance components, and models involving C components, were also tested; see the SOM for details.)

The best fitting model then becomes a 2-factor CP model with an additional pure E-factor (CP2_E1), created by adding an E component to the pure A-factor in the CP1_IP1. This model can be seen as a compromise between the two best fitting models in the initial step: CP1_IP1 and CP3. As the third factor in the CP3 naturally came out as 100% environmental, these two models are almost exactly alike, except that one of the factors in CP1_IP1 is constrained to be 100% genetic (The corresponding CP-factor in CP3 is 69% genetic). CP2_E1 achieves better fit than CP3 only due to having fewer estimated parameters, as the third factor is now constrained to be purely environmental. No further improvements in fit could be achieved from adding or subtracting components for this CP2_E1 model.

Parameter values for CP2_E1 remained similar to those for CP3 and CP1_IP1. The first CP-factor corresponded well with the hypothesized Principled Justice Sensitivity (Principled-JS) trait, having significant positive loadings on all the JS items. The second CP-factor corresponded to Opportunistic Justice Sensitivity (Opportunistic-JS), with positive loadings on Victim items and negative loadings on the rest, with loadings on Victim and Perpetrator items being of larger magnitude than those for Observer and Beneficiary items. Both of these factors were substantially heritable, with heritability-estimates at 45% for the Principled-JS-factor and 69% for the Opportunistic-JS-factor. The purely environmental factor had largely alternating positive- and negative loadings of modest magnitude. Plausibly, this could reflect correlations in measurement errors from how pairs of items are formulated. See figure 4 for an illustration of the model, with all its parameter values. In combination, the three latent factors in the top part of this model explain on average 40% of the variance in each JS-item, ranging from 57% (for item O_2_) to 23% (for item P_2_).

**Figure 4.**
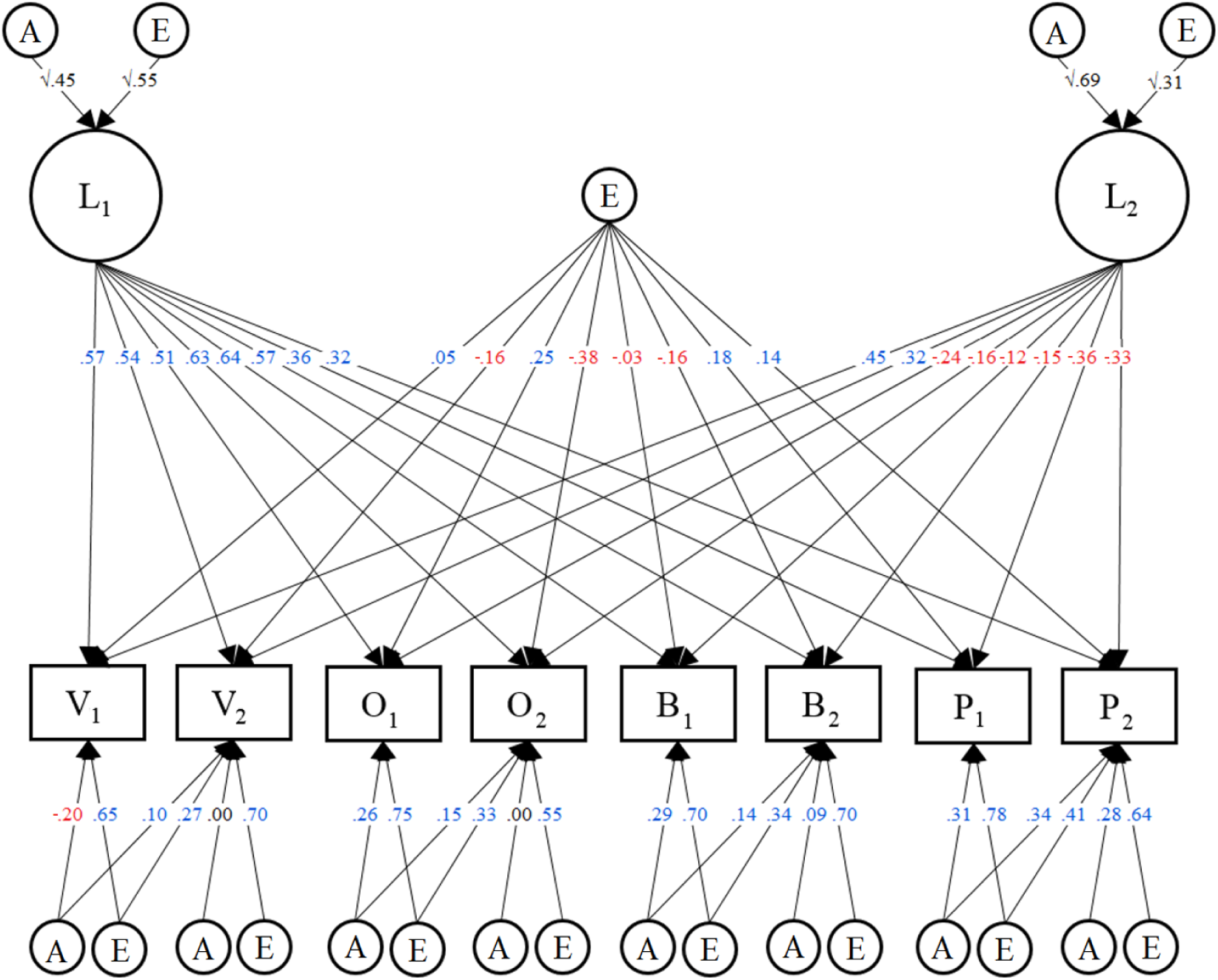
*Note.* Illustration of our best fitting model. Blue numbers are positive and red numbers are negative.

### Correlations of factor scores with relevant variables

We calculated factor scores for the Principled-JS and Opportunistic-JS factors in the best fitting twin model, and correlated these with other theoretically-relevant variables from our dataset (described in the Methods section). See Table 4 for an overview.

**Table 4.** Model comparisons for multivariate twin models.

| Model | EP | -2LL | df | $\Delta$ -2LL | $\Delta$ df | AIC | p |
| --- | --- | --- | --- | --- | --- | --- | --- |
| *Initial model comparisons* |  |  |  |  |  |  |  |
| Cholesky ACE | 116 | 55053.12 | 15353 | NA | NA | 24347.12 | NA |
| Cholesky AE | 80 | 55062.06 | 15389 | 8.940 | 36 | 24284.06 | .999 |
| CP1 | 42 | 55276.22 | 15428 | 223.100 | 75 | 24420.22 | <0.001 |
| CP2 | 52 | 55127.60 | 15419 | 74.475 | 66 | 24289.60 | .222 |
| CP3 | 62 | 55083.79 | 15410 | 30.668 | 57 | 24263.79 | .998 |
| IP1 | 48 | 55190.09 | 15421 | 136.962 | 68 | 24348.09 | <0.001 |
| IP2 | 64 | 55081.24 | 15405 | 28.117 | 52 | 24271.24 | .997 |
| IP3 | 80 | 55066.91 | 15389 | 13.788 | 36 | 24288.91 | .999 |
| **CP1_IP1** | **58** | **55085.92** | **15414** | **32.794** | **61** | **24261.92** | **.998** |
| CP1_IP2 | 74 | 55067.49 | 15400 | 14.364 | 47 | 24267.49 | .999 |
| CP2_IP1 | 68 | 55074.08 | 15405 | 20.961 | 52 | 24264.08 | .999 |
| *Adjustments of best model* |  |  |  |  |  |  |  |
| **CP2_E1** | **60** | **55083.79** | **15412** | **30.668** | **59** | **24259.79** | **.999** |
| CP2_A1 | 60 | 55276.22 | 15428 | 223.100 | 75 | 24420.22 | <0.001 |
| CP1_IP1_A1 | 52 | 55127.60 | 15419 | 74.475 | 66 | 24289.60 | .222 |
| CP1_IP1_E1 | 62 | 55083.79 | 15410 | 30.668 | 57 | 24263.79 | .998 |

*Note.* EP = number of estimated parameters; LL = LogLikelihood; df = degrees of freedom; $\Delta$ = difference as compared to base model; AIC = Akaike's Information Criterion; p = p-values from test of difference from base model. Best fitting model indicated in bold. CP and IP stand for common- and independent pathway respectively. The numbers after CP and IP indicate the number of factors of that kind in the model. E1 signifies the addition of an extra E-factor to a model. All models in the table except the Cholesky ACE are pure AE-models, with no C-components.

Owing in part to our large dataset, most of our correlations were significant, with p-values far below 0.05, even though phenotypic correlations were generally modest in size. The Opportunistic-JS-factor had positive correlations with Social Dominance Orientation, and Neuroticism from the Big Five. It had negative correlations with Openness to Experience and Agreeableness from the Big Five, with Altruism and Interpersonal Trust, and with support for immigration and foreign aid. The Principled-JS-factor correlated positively with Neuroticism and support for immigration and aid, and negatively with Social Dominance Orientation, Agreeableness and Conscientiousness. As mentioned in the methods section, the factors in our model were specified to orthogonal, so Principled- and Opportunistic-JS do not correlate with each other.

Table 4 also contains genetic and environmental correlations (rA and rE, respectively). These describe the degree of overlap in genetic and environmental influences on the pairs of traits. As our genetic correlations are generally higher than the corresponding environmental correlations, the relationships between our latent factors and the other variables seem to be largely due to common genetic influences on both traits, more so than common environmental influences.

**Table 4.** Correlations between factor scores and relevant traits

|  | Principled-JS |  |  | Opportunistic-JS |  |  |
| --- | --- | --- | --- | --- | --- | --- |
|  | rP | rA | rE | rP | rA | rE |
| SDO | **-.10** | **-.13** | -.08 | **.27** | **.40** | **.22** |
| Immi-aid | **.13** | **.25** | .05 | **-.25** | **-.60** | .01 |
| B5O | .02 | .06 | -.01 | **-.16** | **-.28** | -.07 |
| B5C | **-.13** | **-.26** | -.06 | -.03 | **-.10** | -.00 |
| B5E | -.08 | **-.18** | -.01 | -.05 | **-.17** | .05 |
| B5A | **-.12** | **-.14** | **-.10** | **-.17** | **-.19** | **-.16** |
| B5N | **.17** | **.35** | .07 | .08 | **.15** | .03 |
| Altruism | .05 | .05 | .06 | **-.18** | **-.46** | -.01 |
| Trust | -.02 | **-.10** | .02 | **-.20** | **-.43** | -.09 |

*Note.* Principled-JS = factor scores from the factor with significant positive loadings on all 8 JS-items; Opportunistic-JS = factor scores from the factor with positive loadings on the two Victim sensitivity items and negative loadings on the remaining 6 items. rP = Phenotypic correlation; rA = Additive genetic correlation; rE = non-shared environmental correlation. SDO = Social Dominance Orientation; Immi-aid = support for immigration and foreign aid; B5O = Big Five Openness to Experience; B5C = Big Five Conscientiousness; B5E = Big Five Extraversion; B5A = Big Five Agreeableness; B5N = Big Five Neuroticism; Altruism = Frequency of various altruistic behaviors; Trust = Interpersonal Trust. Bold font is used for numbers $>|.10|$ , to highlight practical significance. All correlations that are larger than $|.07|$ are statistically significant at $p < .01$ .

## Discussion

The present study suggests that responses to injustice can be shaped by both principled and self-serving motivations, that these are reflected in latent traits that are genetically-grounded, and that they are genetically associated with other personality and attitudinal measures.

We used genetically-informative data to investigate the existence of latent traits underlying correlations between facets on the Justice Sensitivity scale. In doing this, we avoid the pitfall of factor analyses based purely on phenotypic correlations, which can sometimes support inferring latent psychological traits that do not in fact exist (Franic, 2013). The use of a large twin dataset enabled the construction and comparison of a series of biometric factor models to identify latent factors. We followed the logic that if correlations between facets of a scale are actually mediated by latent psychological traits, then when a factor is split up into separate genetic and environmental components, the patterns of loadings from these components should be highly similar, since they both affect covariances only through their effects on the same latent trait. This prediction is tested through comparing Common and Independent Pathway (CP and IP) models on the Justice Sensitivity (JS) scale, with residuals from JS items belonging to the same facet being allowed to correlate.

We found that the best fitting model had two CP factors and an additional purely environmental factor. One of the CP factors corresponded well with the notion of principled justice sensitivity (Principled-JS), as it had substantial positive loadings on all the JS items, implying that the previously observed association between JS facets is driven by an underlying sensitivity to injustice regardless of the perspective from which it is perceived.

The other CP factor quite strongly resembled the predicted trait of opportunistic justice sensitivity (Opportunistic-JS). Loadings on JS-Victim were substantial and positive, while all other loadings – particularly those for JS-Perpetrator – were negative. This corresponds to a selective sensitivity to injustice that harms one’s interests – a form of moral opportunism. The existence of both principled and opportunistic JS traits explains how the correlation between JS-Victim and JS-Perpetrator is weak even as both of these correlate substantially with the other two JS facets: It results from Victim and Perpetrator sensitivity being pulled in the same direction by Principled-JS, but then also (less strongly) in opposite directions by Opportunistic-JS.

The use of a large twin sample also enabled us to examine heritability in the two justice sensitivity traits. We found that the best fitting models for our data were generally AE models, in which all shared-environmental components were taken out. This suggests that environmental influences that serve to make twins in a pair more similar to each other, such as family environments or socio-economic status in childhood, play little role in explaining variability in principled sensitivity to injustice or in moral opportunism in our sample.

Specifically, we found that 45 % of the variance in Principled-JS was attributable to genes, with the remaining variance being explained by non-shared-environmental influences. This heritability estimate is on a par with those found in the domain of personality (Bouchard & McGue, 2003). The Opportunistic-JS factor was more heritable than the Principled-JS factor, with 69% of its variance observed to be genetic in origin. It should be noted that whenever there are gene-environment correlations, such that genes cause individuals to elicit and/or actively seek out certain environmental influences on a trait, these influences will be interpreted by twin study analyses to be genetic (Purcell, 2002), in line with the notion of an *extended genotype* (Lynch, 2017).

The third, purely non-shared-environmental, factor in our final model had generally weaker loadings than the other two factors, and these were consistently more positive for the first than the second item in each pair of items from the same JS facet. A plausible interpretation of this factor is that it represents correlated measurement errors, related to how pairs of items from the same facet are formulated. The first items from all pairs are formulated quite similarly, just with differences in a few words, and the same is true for all the second items in each pair.

This possibility is consistent with comparisons between the best-fitting biometric model and the results of our phenotypic Confirmatory Factor Analysis on the same data. Here it could be seen that whereas the Principled-JS factor is present in both models, the Opportunistic-JS factor and the added E factor in the twin model are combined into a single factor in the best fitting phenotypic CFA model. Our study thus highlights that, generally, the estimation of separate genetic and environmental factors is helpful in distinguishing methodological artifacts from real psychological traits. By definition, artifacts will tend to not be influenced by genetics or family environments. Real psychological traits, in contrast, will consistently be heritable to some extent (Plomin et al., 2016).

The biometric model results, in addition to the high heritability estimates for both Principled- and Opportunistic-JS, thus speak against the possibility that these factors are methodological artifacts. All models with CP and/or IP factors also include estimates of genetic and environmental influences specific to each item beyond what is accounted for by these broader factors. For all our models, fit was improved through allowing correlations between these influences for items belonging to the same JS facet. This is consistent with prior work on the JS scale, indicating that the current division of items into four separate facets is meaningful. The genetic components of the variance in scores left unexplained by the general factors were mostly shared between items from the same facet, while the non-genetic components of this variance were largely item-specific.

### Construct validation of Principled and Opportunistic Justice Sensitivity

To validate our interpretations of the Principled- and Opportunistic-JS factors, we investigated correlations between their factor scores and other theoretically-relevant variables. Our ability to disentangle motivations for enforcing norms about justice reveals how these motivations differ in their relationships to key personality traits, as well as other measures related to morality, including social dominance orientation, prosociality, and policy support. In addition to phenotypic correlations, we also calculated genetic and environmental correlations, which indicate the degree of overlap in the genetic and environmental factors influencing the traits.

First, we found that the phenotypic correlations between the two JS factors and other traits and attitudes were generally weak, indicating that principled and opportunistic justice sensitivity are moral personality (or ‘character’) traits that are distinguishable from those other constructs in significant ways. This addresses a concern about current personality theories, which is that they do not fully capture how people can vary in moral character (Cawley III, Martin, & Johnson, 2000). While human behavior is often guided by values and moral norms, personality inventories tend mostly to index mere action-tendencies and temperaments, which lack this normative component (Goldie, 2004).

While the two components of justice sensitivity are distinct from personality and social attitudes, we nevertheless found evidence that they have a shared genetic grounding. Generally, we found that genetic correlations were stronger than phenotypic correlations, and that environmental correlations were weak. This indicates that where the two forms of justice sensitivity do overlap with other traits and preferences, they do so because of shared genetic influences on both variables. This corresponds to Cheverud’s conjecture (Cheverud, 1988) that genetic correlations are a representation of the relationship between variables when this is not watered down by measurement errors the way phenotypic correlations can be.

Specifically, we found that principled and opportunistic JS factors had diverging correlations with social dominance orientation (SDO) and support for immigration and foreign aid. Opportunistic-JS had a substantial positive correlation with SDO and policies disfavouring ‘outsiders’, while Principled-JS exhibited the reverse pattern. This accorded with our predictions: Seeking to establish hierarchies from which one benefits can be seen as both opportunistic (high Opportunistic-JS) and unprincipled (low Principled-JS).

Previous research has reported negative correlations of victim sensitivity with facets of agreeableness, trust, and honesty/humility, and positive correlations with neuroticism, jealousy and narcissistic rivalry (see Baumert & Schmitt, 2016). Thus, high victim sensitivity has been variously interpreted as antagonistic self-protection (Back et al., 2013), fear of being exploited (Gollwitzer et al., 2005), and generally as reflecting an antisocial, uncooperative disposition (Baumert & Schmitt, 2016). The present study suggests that these descriptions are perhaps better applied to the broader Opportunistic-JS factor, rather than just to JS-Victim. The negative correlations of Opportunistic-JS with agreeableness, altruism, and trust, and its positive correlations with SDO, suggest that the construct is related to the “dark triad” of personality traits (i.e., Machiavellianism, narcissism, and psychopathy), a possibility worth exploring in future work. It should be mentioned here that our altruism scale (the Self-Report Altruism scale; Rushton, Chrisjohn, & Fekken, 1981) had a low Cronbach’s alpha of .48 in our sample, so our findings should be interpreted with this in mind.

One surprising finding was that of a weakly *negative* correlation between Principled-JS and agreeableness. A possible interpretation of this could be that the desire to uphold justice that comes with having a high score on Principled-JS can sometimes come into conflict with the desire to maintain social harmony associated with agreeableness. Relatedly, it was also surprising that Principled-JS did not have any kind of clear relationship with either altruism nor interpersonal trust. We here predicted substantial positive correlations, but found that both correlations were very close to zero. Our finding is consistent, however, with previous findings that emotional empathy (as contrasted with cognitive empathy) is unrelated to a composite of Observer and Beneficiary sensitivity (Decety & Yoder, 2016), which are the highest loading facets on the Principled-JS factor. Having an empathic and altruistic disposition seems to express itself as low Opportunistic-JS (i.e. lower victim-sensitivity and higher sensitivity to other injustices), rather than simply as high sensitivity to all injustice.

### Limitations and future directions

Although this study goes further than previous analyses of moral traits, it is still limited in what it can be used to infer. We used the short version of the JS scale, which, despite having good construct and criterion validity (e.g., Baumert, et al., 2014; Stavrova, Schlösser, & Baumert, 2014), only directly measured the typical intensity of emotional responses to the various injustice perspectives, leaving out the frequency of injustice responses as well as their typical duration.

Another limitation concerns how our analytic approach was constrained to models with orthogonal factors only, thereby restricting our ability to explore how factors relate to each other. It seems clearly plausible for principled and opportunistc justice sensitivity to correlate negatively, so a more accurate model could be one that allowed for this.

Our sample consists of twin-pairs aged between 60 and 65 years, mostly born and raised in Norway. This limits the generalizability of our findings to younger cohorts and to other cultures. Future work on this topic could seek to replicate our findings in other cultural contexts.

Additionally, our interpretations of the two JS factors can also be validated further. This could be done by including additional theoretically-relevant variables to test the prediction that they will genetically overlap with the heritable traits for principled and opportunistic justice sensitivity we have identified here. For the Opportunistic-JS factor, conceptually-related variables to which we did not have access include the Honesty/Humility-trait in the HEXACO-PI (Ashton & Lee, 2009) and measures of Machiavellianism such as the Machiavellian Personality Scale (Dahling, Whitaker, & Levy, 2009). For Principled-JS, relevant variables include the Fairness dimension in Moral Foundations theory (Graham, Haidt, & Nosek, 2009) and the Morality sub-scale of the Mosher Guilt Inventory (Mosher, 1998).

It would also be important to establish if high principled justice sensitivity or, rather, low moral opportunism is the best predictor of actual altruistic prosocial behavior, including economic behavior in experimental games. We found that self-reports of all variables related to self-sacrificing and cooperative strategies for the greater common good were most strongly related to low moral opportunism, rather than to principled moral impartiality. This suggests that the *absence* of exploitative opportunism may form the core of moral character. Being truly generous and altruistic might imply being *un-principled* about justice, but in the opposite way as a self-serving opportunist would be, so that victim sensitivity is reduced and the aversion to perpetrating injustice increases.

## Conclusion

Evidently, sensitivity to injustice can be driven by both principled- and opportunistic motivations. A principled motivation to preserve justice for its own sake leads to condemnation of all injustice, regardless of the perspective it is perceived from. An opportunistic motivation to use justice in service of other goals rather leads to selective enforcement of rules; namely an *increased* sensitivity to injustices that one is victimized by, combined with a *decreased* tendency to feel shame and guilt when passively benefitting from, or actively perpetrating, injustice towards others.

Our study highlights the complex relationship between justice sensitivity and morality. Moral rules generally have an in-built impartiality, such that they sometimes are beneficial to us and at other times are a hindrance. A principled person reacts in proportion to what the situation demands in moral terms, regardless of who stands to benefit. The moral opportunist, on the other hand, exaggerates or downplays their reactions, depending on what best serves their selfish interests in the current context. For most people most of the time, reactions to injustice will be compromises between both these kinds of motivations.

